# Receptor Activity-Modifying Protein 2 (RAMP2) alters glucagon receptor trafficking in hepatocytes with functional effects on receptor signalling

**DOI:** 10.1101/2021.05.09.443291

**Authors:** Emma Rose McGlone, Yusman Manchanda, Ben Jones, Phil Pickford, Asuka Inoue, David Carling, Stephen R Bloom, Tricia Tan, Alejandra Tomas

## Abstract

**Objectives:** Receptor Activity-Modifying Protein 2 (RAMP2) is a chaperone protein which allosterically binds to and interacts with the glucagon receptor (GCGR). The aims of this study were to investigate the effects of RAMP2 on GCGR trafficking and signalling in the liver, where glucagon is important for carbohydrate and lipid metabolism.

**Methods:** Subcellular localisation of GCGR in the presence and absence of RAMP2 was investigated using confocal microscopy, trafficking assays and radioligand binding assays in human embryonic kidney (HEK293T) and human hepatoma (Huh7) cells. Mouse embryonic fibroblasts (MEFs) lacking Wiskott Aldrich Syndrome protein and scar homologue (WASH) complex were used to investigate the effect of a halt in recycling of internalised proteins on GCGR signalling in the absence of RAMP2. NanoBiT complementation and cyclic AMP assays were used to study the functional effect of RAMP2 on recruitment and activation of GCGR signalling mediators. Response to hepatic RAMP2 up-regulation in lean and obese adult mice using a bespoke adeno-associated viral vector was also studied.

**Results:** GCGR is predominantly localised at the plasma membrane in the absence of RAMP2 and exhibits remarkably slow internalisation in response to agonist stimulation. Rapid intracellular retention of glucagon-stimulated GCGR in cells lacking WASH complex indicates that activated GCGRs undergo continuous cycles of internalisation and recycling despite apparent GCGR plasma membrane localisation up to 40 minutes post-stimulation. Co-expression of RAMP2 induces GCGR internalisation both basally and in response to agonist-stimulation. The intracellular retention of GCGR in the presence of RAMP2 confers a bias away from β-arrestin-2 recruitment coupled to increased activation of G_αs_ proteins at endosomes. This is associated with increased short-term efficacy for glucagon-stimulated cAMP production, although long-term signalling is dampened by increased receptor lysosomal targeting for degradation. Despite these signalling effects, only minor disturbance of carbohydrate metabolism was observed in mice with up-regulated hepatic RAMP2.

**Conclusions:** By retaining GCGR intracellularly, RAMP2 alters the spatiotemporal pattern of GCGR signalling. Further exploration of the effects of RAMP2 on GCGR *in vivo* is warranted.

**Graphical abstract:** Icons sourced from [1]

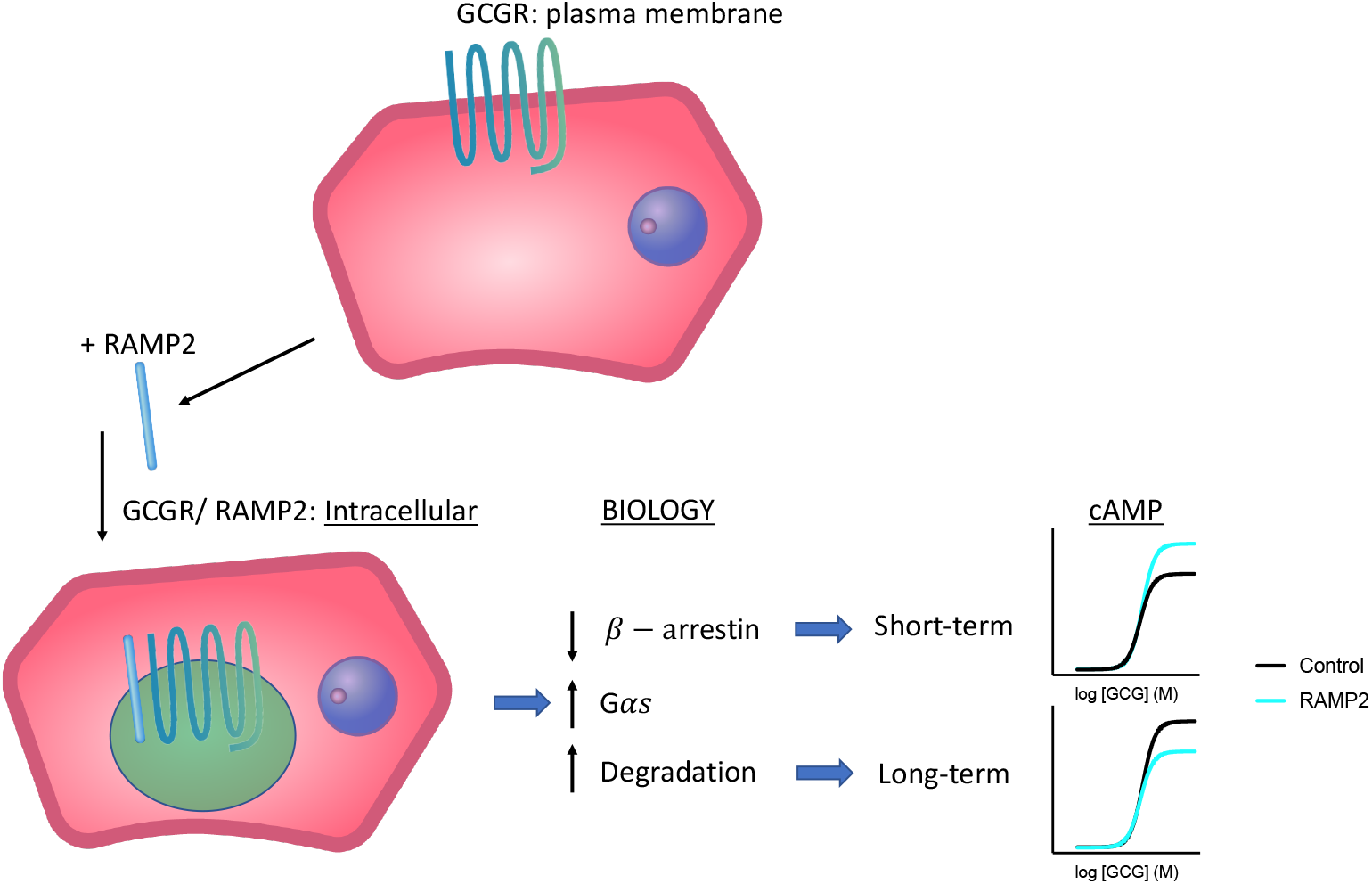

## 1 Introduction

Glucagon acts through the glucagon receptor (GCGR), a prototypical G protein-coupled receptor (GPCR) of the secretin-like (class B) family [2]. The effects of glucagon on the liver include increased hepatic glucose output by stimulation of glycogenolysis and gluconeogenesis, inhibition of *de novo* lipogenesis and increased fatty acid oxidation [3–5]. Type 2 diabetes mellitus (T2DM) and non-alcoholic fatty liver disease (NAFLD) are characterised by high glucagon levels and glucagon resistance [6; 7]: manipulation of glucagon signalling is a potential pharmacological strategy for treatment of these conditions [8; 9]. Intracellular trafficking of other GPCRs from the secretin-like family has been demonstrated to play a key role in the regulation of receptor signalling outputs [10–14], suggesting that modifying intracellular trafficking may be a tractable approach to modulate GCGR signalling.

Receptor-activity modifying proteins (RAMPs) are mammalian accessory proteins that interact allosterically with the vast majority of GPCRs [15; 16]. Their actions are wide-reaching and highly variable but include modulation of receptor trafficking, changes to ligand specificity and alteration of intracellular response to receptor activation [17]. As RAMPs interact with their cognate GPCRs in a complex lipid membrane environment, their effect on receptor function varies depending on the cell type in which they are studied [18]. We, and others, have previously demonstrated that RAMP2 (but not RAMP1 or RAMP3) co-localises with the GCGR and alters its pharmacology in certain cell types [19–21]. We also observed a reduction in cell surface GCGR in the presence of RAMP2 [20]. The aims of the present study were twofold: firstly, to analyse the effect of RAMP2 on the intracellular trafficking and spatiotemporal regulation of signalling of the GCGR in more detail; and secondly, to investigate the effects of the interaction between RAMP2 and the GCGR in hepatocytes, a physiologically relevant cell type.

## 2 Materials and Methods

Please also see Supplementary Methods.

### 2.1 Peptides

All peptides were purchased from Insight Biotechnology. Glucagon(1-39) and glucagon-like peptide 1 (GLP-1) (7-36)NH_2_, the predominant bioactive forms of glucagon and GLP-1, respectively, were used for all experiments, except for some instances in which fluorescent glucagon and GLP-1 peptide conjugates featuring a fluorescein isothiocyanate (FITC-GCG and FITC-GLP-1) were used to monitor ligand binding and/or uptake. FITC-GLP-1 has been previously described and validated [22]. For radioligand binding assays, glucagon was directly iodinated in-house (I^125^ from Hartmann Analytic) and purified with reversed-phase high performance liquid chromatography [23].

### 2.2 Cell lines

HEK293T, MEF (flox/flox and WASH-out, a gift from Professor Daniel Billadeau, Mayo Clinic, Rochester, USA) and Huh7 hepatoma cells were maintained in DMEM supplemented with 10% FBS and 1% penicillin/streptomycin and cultured at 37°C in a 5% CO_2_ atmosphere. INS-1 832/3 cells, a gift from Professor Christopher Newgard, Duke University Medical Centre, Durham, USA, were maintained in RPMI supplemented with 11 mM glucose, 10% FBS, 10 mM HEPES, 2 mM L-glutamine, 1 mM pyruvate, 50 μM β-mercaptoethanol and 1% penicillin/streptomycin. A stable clone of Huh7 cells expressing human GCGR (Huh7-GCGR) was generated from a previously described multi-clonal cell population [20] by flow cytometric sorting of cells labelled with FITC-GCG, and subsequently maintained in DMEM, 10% FBS, 1% penicillin/streptomycin and 1 mg/ml G418 (Thermo Fisher).

### 2.3 Transfections

Transient transfections of SNAP-GCGR, SNAP-GLP-1R (both Cisbio), RAMP2, GCGR-GFP (both Origene), RAMP2-CFP, empty vector (EV)-CFP (both GeneCopoeia), Nb37-GFP (a gift from Professor Roshanak Irannejad, University of California San Francisco, USA), CLIP-RAMP2 [cloned in-house and sequence-verified from RAMP2 (Origene) and CLIP-β2-AR (a gift from Professor Davide Calebiro, University of Birmingham)], TGN-marker (Venus-tagged GRIP domain, made in-house), GLP-1R-GFP (a gift from Professor Alessandro Bisello, University of Pittsburgh, USA), HALO-GCGR and HALO-GLP-1R (both made in-house), Rab5-Venus (a gift from Professor Kevin Pfleger, University of Western Australia) and plasmids for the NanoBiT complementation assays (see Supplementary Methods) were performed using Lipofectamine 2000 (Thermo Fisher) for HEK293T and Huh7 cells, or by electroporation with the Neon transfection system (Thermo Fisher) for MEF cells, according to the manufacturer’s instructions. Experiments were performed 24 hours after transfection unless otherwise indicated. Reverse transfection with siRNA against *RAMP2* or a Silencer Select negative control (both Ambion), again with Lipofectamine 2000, was used to downregulate RAMP2. Reagents were added at the time of plating the cells, and experiments performed 72 hours later.

### 2.4 Antibodies

SNAP-GCGR was detected with an anti-SNAP-tag rabbit polyclonal antibody (P9310S, New England Biolabs, 1/500) followed by goat anti-rabbit IgG H&L HRP (ab6271, Abcam, 1/2,000). Post stripping, tubulin was labelled with anti-α-tubulin mouse monoclonal antibody (T5168, Sigma, 1/5,000) followed by sheep anti-mouse secondary antibody HRP (ab6721, Abcam, 1/5,000). For liver samples, the following antibodies were used: anti-RAMP2 sc-365240 at 1/500 dilution; secondary sc-516102 at 1/1,000 (both from Santa Cruz Biotechnology); and anti-GAPDH mab374 at 1/500 dilution (Merck); secondary #15014 at 1/10,000 (Active Motif).

### 2.5 Animal care

Experiments were performed in accordance with the UK Animals (Scientific Procedures) Act 1986 and approved by the Animal Welfare and Ethical Review Board at Imperial College London. C57BL/6J male mice (Charles River) were group housed in cages at controlled temperature (22°C), and a 12-hour light-dark cycle with free access to water. All interventions were performed during the light cycle. Mice were weaned and maintained on standard chow (11% kcal from fat and 62% from carbohydrate, SDS Rm3).

### 2.6 Up-regulation of hepatic RAMP2 in mice

Mouse *Ramp2* and *GFP* (control) under the albumin promoter were constructed in an AAV2/8 pseudotyped adeno-associated virus vector (Vector BioLabs). At age 6 weeks, mice were administered a tail vein intravenous injection of 1×10^11^ gene count of AAV-alb-*GFP* or AAV-alb-*Ramp2*. Mice were randomised for injections and returned to their original cage: that is, mice with hepatic RAMP2 upregulation were co-housed with control mice. After 2-3 weeks, metabolic tests on lean mice were performed. At age 13 weeks mice were transferred to a high fat diet containing 60% kcal from fat (Research Diets D12492). After a further 8 weeks, metabolic tests were performed on obese mice. All tests were performed in 5-hour fasted mice unless otherwise specified. Tail vein blood glucose was measured using a handheld glucometer (Nexus, GlucoRx) before and at indicated intervals after intraperitoneal injections of glucose (2 mg/kg body weight), insulin (0.5 or 1 U/kg of Actrapid human insulin for lean and obese mice respectively, Novo Nordisk), pyruvate (2g/kg, Sigma) or glucagon (10 nmol/kg body weight). Obese mice were dosed with glucose and glucagon according to estimated lean weight of the same strain, sex and age of mouse maintained on standard chow (31 g). After the study period, mice were culled via decapitation following a 5-hour period of food restriction. The liver was harvested rapidly, and snap frozen in liquid nitrogen. Hepatocytes were isolated using a collagenase perfusion, as previously described [24]. After washing and plating, they were serum-starved overnight before cyclic adenosine monophosphate (cAMP) assays were performed.

### 2.7 Data and statistical analyses

All statistical analysis was performed using GraphPad Prism 8.0. For cAMP and NanoBiT complementation assays, E_max_ and logEC_50_ were derived for each repeat and then compared using paired t-tests. Manders’ coefficient was calculated by comparing the confocal images in Image J using the Coloc2 plugin to illustrate the extent of co-localisation between two fluorophore markers [25]. Signal bias calculations were derived from NanoBiT data. Baseline-corrected curves, normalised to vehicle, were generated for glucagon-stimulated LgBiT-mini-G_s_, LgBiT-mini-G_q_ and LgBiT-β-arrestin-2 recruitment data to GCGR-SmBiT in both RAMP2- and pcDNA3.1-transfected HEK293T cells and used to calculate area under the curve (AUC) over 30 minutes (see Supplementary Methods). RAMP2/pcDNA3.1 AUC ratios were subsequently calculated for each recruited factor and compared for statistical significance with a one-way ANOVA with Dunnett’s post-hoc test. All other specific statistical tests are indicated in the figure legends. AUC was calculated from y=0. Statistical significance is considered with p<0.05.

## 3 Results

### 3.1 GCGR exhibits rapid internalisation and recycling to the cell membrane upon agonist stimulation

Unlike other glucagon-like peptide receptors, GCGR does not appear to exhibit short-term agonist-stimulated internalisation [13; 26; 27]. To investigate this phenomenon, we used a fluorescent ligand (FITC-GCG) to stimulate SNAP-GCGR-expressing HEK293T cells, which were chosen because they do not express endogenous RAMPs [21]. FITC-GCG has comparable potency for cAMP production to glucagon at the GCGR (logEC_50_ −8.8 vs −9.1; p=0.11, Supplementary Figure 1A). Although FITC-GCG rapidly accumulated inside the cell after only a few minutes of stimulation, internalisation of the GCGR occurred much more slowly (Figure 1A and B). This is in stark contrast to the rapid internalisation exhibited by both FITC-GLP-1 and the glucagon-like peptide 1 receptor (GLP-1R) (Figure 1C and D), in agreement with previous observations indicating that, following agonist stimulation, the GLP-1R is rapidly internalised within 10 to 15 minutes of GLP-1 exposure [12; 13; 26]. Similar findings of minimal GCGR internalisation in contrast to substantial GLP-1R internalisation were observed in a pancreatic beta cell line after 30 minutes of agonist stimulation (Supplementary Figure 1B and C); and in HEK293T cells using receptors with a C-terminal GFP tag and unlabelled agonists (Supplementary Figure 1D and E).

**Figure 1 –.**
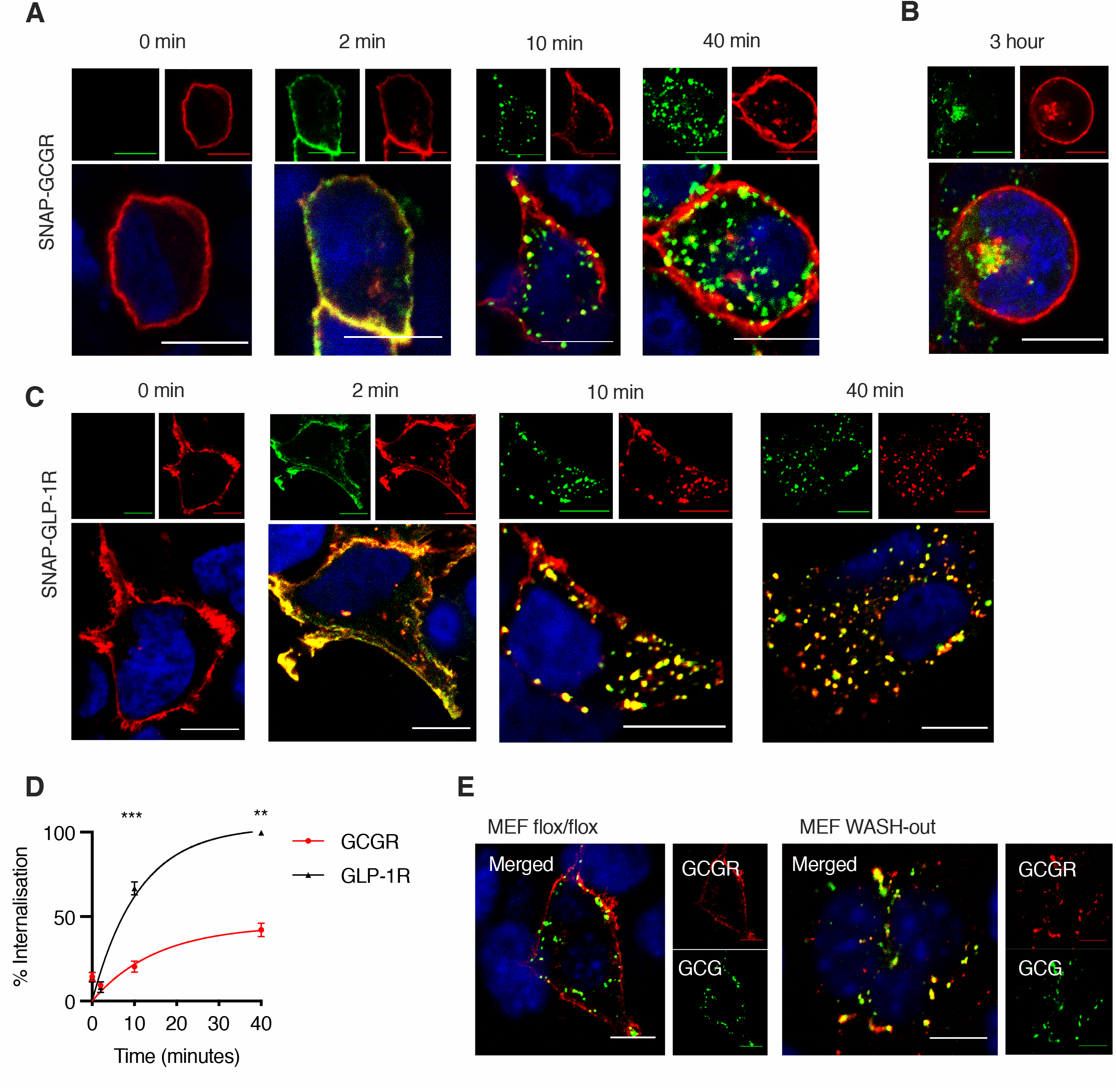
GCGR rapidly recycles to the plasma membrane following glucagon stimulation. A-C: HEK293T cells transfected with SNAP-GCGR (A, B) or SNAP-GLP-1R (C), labelled with SNAP-Surface 549 (red), and stimulated with 100 nM FITC-GCG or FITC-GLP-1 (green), respectively, for the indicated time periods; nuclei stained with DAPI (blue); scale bars = 10 μm. D: Percentage of internalisation of SNAP-GCGR vs. SNAP-GLP-1R in response to, respectively, FITC-GCG or FITC-GLP-1, mean ± SEM of n= 4 pooled data shown, fitted to exponential plateau and analysed using mixed effects model with Sidak’s multiple comparison test; **p<0.01, ***p<0.001. E: MEF cells with or without WASH knockout (MEF flox/flox control or WASH-out), transfected with SNAP-GCGR (labelled with SNAP-Surface 549, red) and stimulated with FITC-GCG (green) for 30 minutes; nuclei stained with DAPI (blue); scale bars = 10 μm.

Given the discrepancy between internalisation of the GCGR and its ligand, we hypothesised that the apparent lack of GCGR internalisation is illusory: for shorter stimulation periods the GCGR would internalise along with its ligand, which it would deposit intracellularly, but then it would rapidly return to the cell membrane by a fast-recycling pathway, further undergoing rapid cycles of internalisation and recycling before a substantial proportion of the receptor would accumulate intracellularly. To investigate this hypothesis, we employed mouse embryonic fibroblasts (MEFs) in which the Wiskott-Aldrich syndrome protein and SCAR homologue (WASH) complex has been knocked out [28]. WASH is an important regulator of vesicle recycling in many cell types, whose deficiency non-specifically traps internalised GPCRs in endosomal compartments [28–31]. Following a short period of glucagon stimulation, we found the GCGR was retained intracellularly in MEFs lacking WASH (WASH-out MEFs), while the receptor localised primarily at the plasma membrane in control MEF flox/flox cells (Figure 1E). These results indicate that, in the absence of RAMP2, GCGR undergoes a continuous cycle of internalisation followed by intracellular ligand deposition and rapid plasma membrane recycling which leads to a slow course of intracellular GCGR accumulation.

### 3.2 The GCGR accumulates intracellularly in the presence of RAMP2

In other contexts, RAMPs have been demonstrated to influence post-endocytic receptor trafficking [32; 33]. To investigate whether RAMP2 affects the subcellular localisation of the GCGR, we cotransfected HEK293T cells with SNAP-GCGR and CLIP-RAMP2, a RAMP2 derivative with a short N-terminal CLIP-tag, and labelled the tagged proteins with fluorescently-conjugated membrane-impermeable surface SNAP-tag and CLIP-tag probes. In the absence of RAMP2, SNAP-GCGR was observed predominately at the cell membrane; in cells expressing RAMP2, however, a proportion of SNAP-GCGR was visible inside the cell where it colocalised with CLIP-RAMP2 (Manders’ coefficient 0.45) (Figure 2A). Similar results were obtained with HALO-tagged GCGR (Supplementary Figure 2A). Co-transfection with Rab5-Venus, an endosomal marker, indicated that SNAP-GCGR and CLIP-RAMP2 co-localise in early endosomes (Figure 2B). Given the physiological relevance of hepatocytes for glucagon signalling and the cell line dependence of RAMP activity, we next examined this phenomenon in Huh7 hepatoma cells, which express low levels of endogenous RAMP2 [20]. Huh7 cells were similarly transfected with SNAP-GCGR with or without untagged RAMP2 co-expression, and surface GCGR labelled with a fluorescent surface SNAP-tag probe. Less SNAP-GCGR was observed at the cell surface with exogenous RAMP2 (Figure 2C and D; p=0.03). To further investigate this phenomenon using an untagged GCGR, we performed a radioligand binding assay to measure GCGR cell surface density in Huh7 cells stably expressing GCGR (Huh7-GCGR) at 4°C (to inhibit receptor endocytosis) where RAMP2 was either up- or down-regulated. We observed a reduced apparent GCGR cell surface density where RAMP2 was up-regulated (Figure 2E and F), while surface GCGR was increased where RAMP2 was down-regulated (Figure 2G; p=0.0096).

**Figure 2 –.**
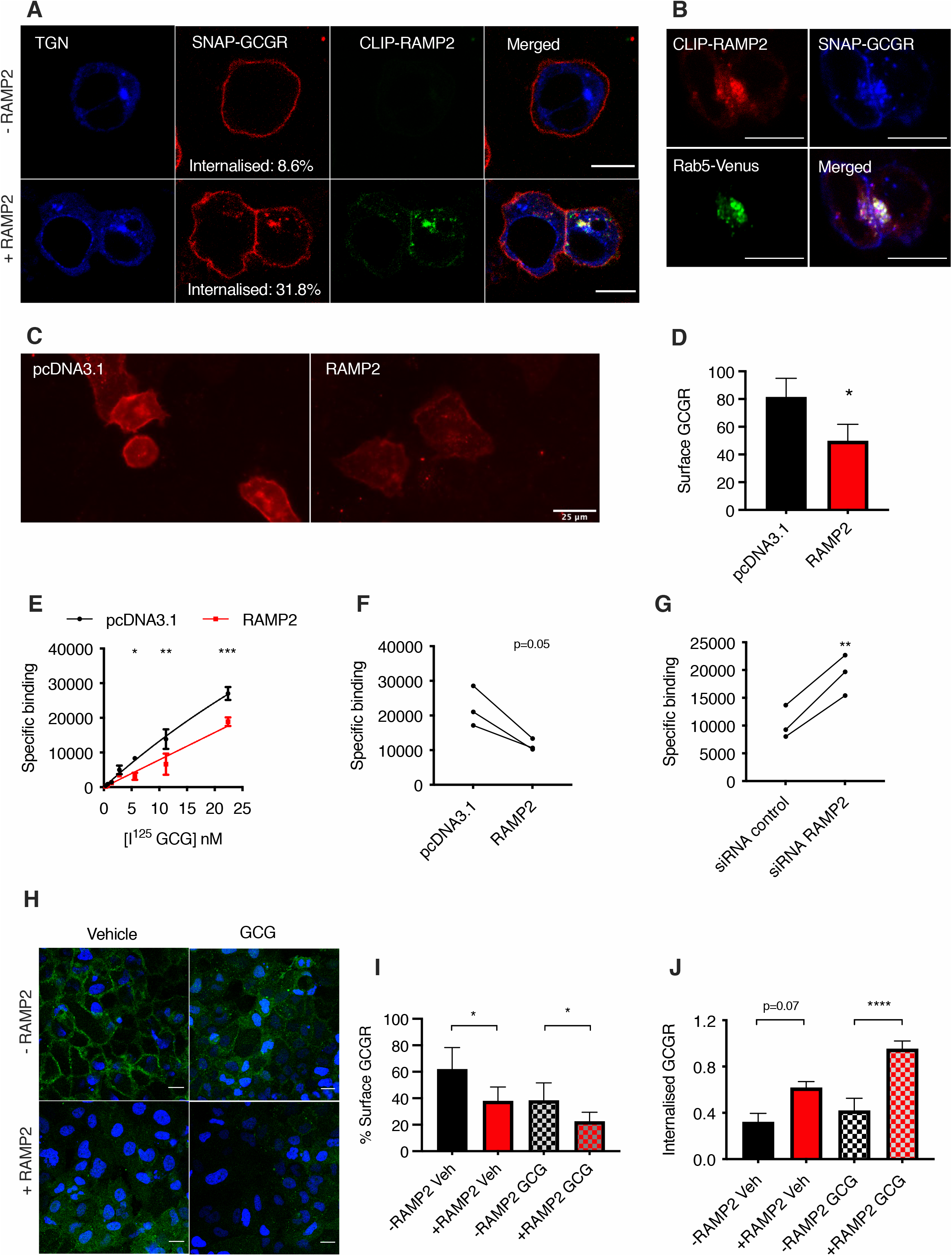
RAMP2 accelerates internalisation of GCGR. A: HEK293T cells transfected with a trans-Golgi network (TGN) marker (blue), SNAP-GCGR (labelled with SNAP-Surface 649, red) and with or without CLIP-RAMP2 (labelled with CLIP-Surface 547, green); scale bars = 10 μm. B: HEK293T cells transfected with Rab5-Venus (green), SNAP-GCGR (labelled with SNAP-Surface 649, blue) and CLIP-RAMP2 (labelled with CLIP-Surface 547, red) and then treated with 100 nM glucagon (GCG) for 40 minutes; scale bars = 10 μm. C: Huh7 cells co-transfected with SNAP-GCGR (labelled with SNAP-Surface 649, red) and CLIP-RAMP2 or control empty vector (pcDNA3.1); scale bars = 25 μm. D: Surface SNAP-GCGR density in unstimulated Huh7 cells from C; *p<0.05. E-G: Specific binding of I^125^-GCG to Huh7-GCGR: increasing I^125^-GCG concentrations in control (pcDNA3.1) *vs*. RAMP2 over-expression (E), and at a single concentration of 5.6 nM I^125^-GCG in control (pcDNA3.1) *vs*. RAMP2 up-regulation (F), or control *vs*. RAMP2 silencing (G), measured in counts over 240 seconds; data are mean ± SEM of n = 3-4 experiments, normalised to total protein levels; *p<0.05; **p<0.01; ***p<0.001. H: Huh7-GCGR cells transfected with RAMP2 and then treated with either vehicle or 100 nM GCG for 30 minutes at 37°C prior to addition of FITC-GCG (green) at 4°C; nuclei stained with DAPI (blue); scale bars = 10 μm. I: Quantification of surface GCGR by FITC-GCG binding in cells from H; *p<0.05; J: HEK293T cells transfected with SNAP-GCGR and either empty vector (EV)-CFP (-RAMP2) or RAMP2-CFP (+RAMP2) and then stimulated with vehicle or 100 nM GCG, SNAP-GCGR seen inside the cell expressed as ratio of total cellular GCGR; ***p<0.0001. Data are mean ± SEM of at least n = 3 independent experiments. Statistical significance was analysed using paired t-tests (D-I) and one-way ANOVA with Sidak’s post-hoc test (J).

We repeated the assay using FITC-GCG at 4°C to quantify the level of surface GCGR in Huh7-GCGR cells with and without RAMP2 co-expression under basal conditions and after GCG stimulation. We found that the presence of RAMP2 decreased plasma membrane GCGR under both conditions (Figure 2H and I). Additionally, we measured an increase in the internalisation propensity of the GCGR in the presence of RAMP2 in HEK293T cells with both the SNAP-tagged (Figure 2J) and the HALO-tagged GCGR (Supplementary Figure 2A and B). Taken together, these findings demonstrate that the presence of RAMP2 increases the intracellular localisation of GCGR, either by increasing the rate of GCGR endocytosis or by reducing the rate of GCGR recycling to the plasma membrane, in both basal and glucagon-stimulated conditions, a consistent finding in different cell lines.

### 3.3 Up-regulation of RAMP2 increases cAMP production acutely and engenders signalling bias in hepatoma cells

Agonist-stimulation of GCGR leads to recruitment of Gα_s_, Gα_i_ and Gα_q_ proteins, triggering intracellular events that lead to modulation of cAMP levels, as well as recruitment of the β-arrestins (Supplementary Figure 3A and B). It has been previously demonstrated that glucagon-stimulated cAMP production is increased in the presence of RAMP2 in HEK293T [21] and Chinese hamster ovary (CHO) cells [20]. Here, we determine that in Huh7-GCGR hepatoma cells, up-regulation of RAMP2 also acutely increases efficacy for cAMP production in response to glucagon stimulation (Figure 3A; Table 1); this is consistent with our previous finding that a reduction of RAMP2 in Huh7 cells is associated with a decrease in cAMP production with glucagon [20]. One possible mechanism explaining this finding could be differential recruitment of intracellular mediators to the GCGR in the presence of RAMP2; indeed, previous work indicated that RAMP2 reduces recruitment of the inhibitory G protein subunit Gα_i_ to GCGR in yeast strains expressing G protein chimeras [21]. To further explore the effect of RAMP2 on the recruitment of G proteins and β-arrestins to the GCGR, we employed a NanoBiT complementation assay in HEK293T cells, which we have previously used to assess transducer coupling by the glucagon family of receptors [26; 34]. We observed a reduction in β-arrestin-2 recruitment to the GCGR in the presence of RAMP2, with no difference in recruitment of mini-G_s_, -G_i_ and -G_q_ proteins (Figure 3B-E), leading to a bias away from β-arrestin-2 recruitment when compared to the recruitment of all three mini-G subunits when RAMP2 is up-regulated (Figure 3F).

**Figure 3 –.**
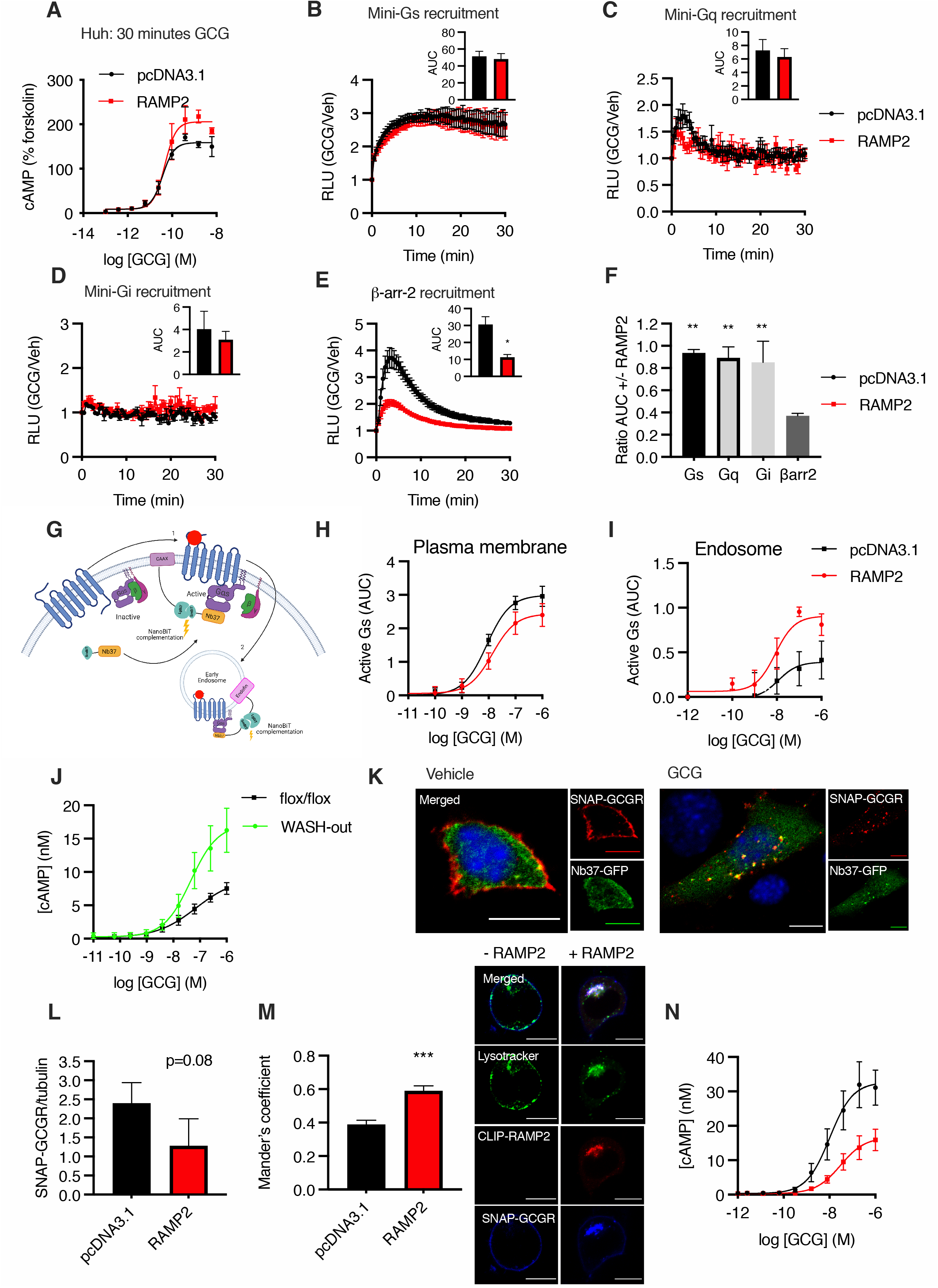
Up-regulation of RAMP2 has spatiotemporal effects on intracellular signalling. A: cAMP dose response curves in Huh7-GCGR cells transfected with RAMP2 or control (pcDNA3.1) after 30 minutes stimulation with GCG, relative to forskolin (10 μM) responses; n = 4; 4-parameter fit of pooled data shown. B-E: Time-course of miniG_s_-, miniG_q_-, miniG_i_- and β-arrestin-2-LgBiT recruitment to GCGR-SmBiT following 100 nM GCG stimulation measured by NanoBiT assays in the presence or absence of RAMP2, with AUC shown as insets. Data normalised to baseline (unstimulated) signal and expressed as Relative Light Units (RLU) as mean ± SEM of n = 4 experiments; *p<0.05. F: Ratio of AUCs in the presence *vs*. absence of RAMP2 for miniG_s_-, miniG_q_-, miniG_i_- and β-arrestin-2-LgBiT recruitment to GCGR-SmBiT; data from B-E with each mini-G protein compared to β-arrestin-2 recruitment; **p<0.01. G: Schematic of NanoBiT subcellular G protein activation assay: the GCGR and heterotrimeric G proteins are quiescent at the cell membrane. When glucagon binds to its receptor, recruited Gα_s_ is activated and binds to Nb37 (1). Nb37-SmBiT complements CAAX-LgBiT upon plasma membrane activation. Following receptor internalisation (2), Nb37-SmBiT complements with Endofin-LgBiT indicating endosomal signalling. Created with BioRender.com. H and I: Gα_s_ activation at the plasma membrane and endosomal membranes, respectively; AUC over 30 minutes normalised to baseline; n = 6; 4-parameter fit of pooled data shown. J: cAMP dose responses to glucagon in control vs. WASH-out MEFs; n = 4; 4-parameter fit of pooled data shown. K: MEF WASH-out cells cotransfected with SNAP-GCGR (labelled with SNAP-Surface 549, red) and Nb37-GFP (green) and stimulated with GCG for 30 minutes: examples of Nb37-GFP co-localised with internalised SNAP-GCGR indicating that it is actively signalling via *a* α_s_ are arrowed; nuclei stained with DAPI (blue); scale bars = 10 μm. L: Western blot quantification of total SNAP-GCGR levels in HEK293T cells with and without RAMP2 co-expression following stimulation with 100 nM GCG for 4 hours, normalised to tubulin as a loading control; data is mean ± SEM of n = 3 repeats (see Supplementary Figure 3E for representative blots). M: HEK293T cells transfected with SNAP-GCGR (labelled with SNAP-Surface 649, blue) with or without CLIP-RAMP2 co-expression (labelled with CLIP-Surface 547, red), and then treated with 100 nM GCG for 3 hours before 5 minutes labelling with LysoTracker Green (green); scale bars = 10 μm; Mander’s coefficient (SNAP-GCGR over lysotracker) quantified; n = 5; ***p<0.001. N: cAMP dose responses to GCG in Huh7-GCGR cells transfected with RAMP2 or control (pcDNA3.1) after a 24-hour stimulation period; n = 4; 4-parameter fit of pooled data shown. Statistical significance analysed using paired t-test (panels B-E, L and M) or one-way ANOVA with Sidak’s post-hoc test (F).

**Table 1 –.** Responses to glucagon stimulation in different cell types for the indicated times. Parameter estimates ± SEM from responses depicted in Figure 3, n = 4-6, *p<0.05, paired t-test.

|  | Huh7-GCGR acute (30 minute) cAMP |  |
| --- | --- | --- |
|  | -RAMP2 | +RAMP2 |
| **Emax** | 141.2 ± 8.6 | 201.6 ± 9.6* |
| **LogEC50** | -10.40 ± 0.14 | -10.21 ± 0.20 |
|  | MEF flox/flox cAMP | MEF WASH-out cAMP |
| **Emax** | 8.49 ± 1.14 | 18.3 ± 2.8* |
| **LogEC50** | -7.33 ± 0.25 | -7.17 ± 0.31 |
|  | HEK293T plasma membrane GCGR activity (NanoBiT) |  |
|  | -RAMP2 | +RAMP2 |
| **Emax** | 3.01 ± 0.26 | 2.48 ± 0.32 |
| **LogEC50** | -8.05 ± 0.06 | -7.81 ± 0.15 |
|  | HEK293T endosomal membrane GCGR activity (NanoBiT) |  |
|  | -RAMP2 | +RAMP2 |
| **Emax** | 0.49 ± 0.15 | 0.92 ± 0.08* |
| **LogEC50** | -7.50 ± 0.35 | -8.24 ± 0.24* |
|  | Huh7-GCGR prolonged (24 hour) cAMP |  |
|  | -RAMP2 | +RAMP2 |
| **Emax** | 34.6 ± 5.4 | 16.4 ± 3.2* |
| **LogEC50** | -7.87 ± 0.26 | -7.59 ± 0.16 |
|  | Primary hepatocytes cAMP |  |
|  | -RAMP2 | +RAMP2 |
| **Emax** | 458 ± 146 | 521 ± 84 |
| **LogEC50** | -10.0 ± 0.81 | -10.5 ± 0.49 |

### 3.4 Endosomal retention of GCGR increases agonist-stimulated cAMP production

Increased intracellular retention of GCGR in the presence of RAMP2 could potentiate cAMP accumulation, as sustained signalling from endosomes has been observed in several related secretin-family GPCRs and has been linked to the formation of GPCR-G protein megacomplexes [30–33]. To investigate this hypothesis, we designed a novel NanoBiT subcellular G protein activation assay to distinguish agonist-stimulated activation of the GCGR at the plasma *versus* endosomal membranes. Here, we co-express a plasma membrane (CAAX) or endosomal (Endofin) marker fused to the large BiT (LgBiT) subunit of the Nanoluc luciferase together with nanobody-37 (Nb37), a single domain antibody which binds specifically to nucleotide-free Gα_s_ in complex with active receptors [35], fused to a complementary small subunit (SmBiT) of Nanoluc. Under this configuration, when the two nanoluciferase subunits are closely apposed, a quantitative luminescent signal is generated [36], indicating the presence of active Gα_s_ in endosomal or plasma membrane compartments (see Figure 3G for a schematic of the assay). In HEK293T cells in the absence of glucagon stimulation, we observed a trend towards diminished basal levels of activation at the plasma membrane in the presence of RAMP2 (p=0.08) but not at the endosomal compartment (Supplementary Figure 3C and D), in keeping with our previous finding of reduced surface GCGR levels in basal conditions. Following baseline normalisation, we observed no difference in activation of plasma membrane GCGR upon glucagon stimulation in the presence of RAMP2 (Figure 3H; Table 1), but we recorded a significant increase in both efficacy and potency for endosomal GCGR signalling with RAMP2 (Figure 3I; Table 1). Furthermore, by artificially inducing GCGR intracellular retention in the MEFs devoid of WASH complex (Figure 1D), we similarly increased the efficacy for cAMP production after glucagon stimulation (Figure 3J, Table 1). Moreover, upon glucagon stimulation in these MEF WASH-out cells, we could observe colocalisation within intracellular puncta of a GFP fusion of Nb37 and the SNAP-GCGR, indicating that the GCGR is active and signalling at this intracellular location (Figure 3K).

Since a proportion of intracellular GPCRs can be targeted from endosomes towards the degradative pathway [37], we next asked if the propensity for degradation of the GCGR might be increased by RAMP2 co-expression. We observed that prolonged (4 hours) glucagon stimulation in the presence of RAMP2 resulted in a trend towards reduced total SNAP-GCGR levels (p=0.08; Figure 3L and Supplementary figure 3E). This was associated with greater co-localisation of SNAP-GCGR with low pH endosomal compartments marked by LysoTracker in the presence of RAMP2 after 3 hours of glucagon stimulation (Figure 3M). We also investigated how cAMP accumulation would be affected by RAMP2 co-expression in Huh7-GCGR cells in the context of prolonged glucagon stimulation and found that, although RAMP2 was associated with increased efficacy for cAMP generation when measured acutely (Figure 3A), this effect was reversed after 24 hours of glucagon exposure (Figure 3N and Table 1).

### 3.5 Hepatic RAMP2 up-regulation does not grossly affect carbohydrate metabolism in lean or obese adult male mice

To investigate whether there was a biological effect of the observed changes in glucagon-stimulated cAMP accumulation in hepatocytes following up-regulation of RAMP2, we used an adeno-associated virus vector to up-regulate murine *Ramp2* gene under the control of the albumin promoter (AAV-alb-RAMP2) in hepatocytes of adult male mice. We confirmed up-regulation of mRNA and protein expression of RAMP2 in the livers of treated mice for at least 4 months post-injection (Supplementary Figure 4A-C). Lean mice treated with AAV-alb-RAMP2 had no readily apparent phenotypic differences, with comparable body weight to mice injected with a control AAV (AAV-alb-GFP) (Figure 4A). Although they exhibited a small reduction in glucose excursion during a glucose tolerance test following a 5hour fast (Figure 4B), the same was not observed after a 24-hour fast (Figure 4C). They also exhibited no significant differences in glycaemic responses when subjected to a glucagon challenge or insulin tolerance test (Figure 4D and E). Since compensatory mechanisms could mask differences in glucagon signalling between the two cohorts, we harvested primary hepatocytes and measured cAMP in response to glucagon: although there was a trend for higher cAMP efficacy and potency in AAV-alb-RAMP2 hepatocytes, this was not statistically significant (Figure 4F and Table 1).

**Figure 4 –.**
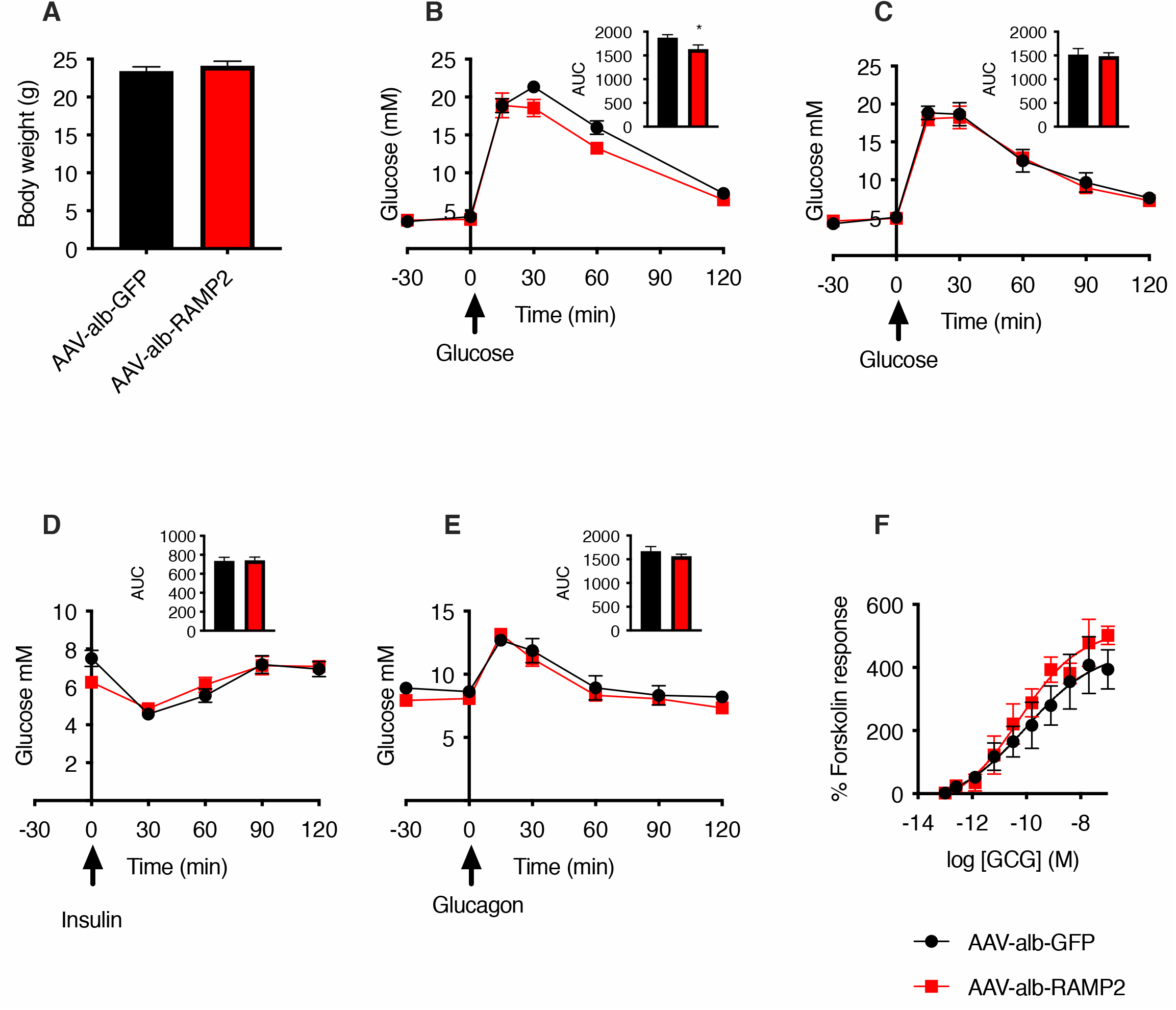
Up-regulation of hepatic RAMP2 in lean adult male mice is not associated with a phenotypic change. A: Mouse body weight after 18 days up-regulation of hepatic mouse RAMP2 (AAV-alb-RAMP2) or control (AAV-alb-GFP); n = 4; data are mean ± SEM. B: Glucose tolerance test after 5-hour fast. C: Glucose tolerance test after 24-hour fast. D: Insulin tolerance test. E: Glucagon challenge. B-E experiments performed in mice 3 to 4 weeks post-AAV injection; n = 9-10 per group; data are mean ± SEM, with AUC shown as inset; unpaired t-test; *p<0.05. F: cAMP dose response to GCG in isolated primary hepatocytes from AAV-alb-GFP and AAV-alb-RAMP2 mice, normalised to forskolin (10 μM) responses; n = 4; 4-parameter fit of pooled data shown; n = 7 separate mice from each cohort harvested over 4 days.

The same cohort of mice were next transferred to a high-fat diet, and metabolic tests were performed when the mice were obese: again, there were no apparent differences between the groups (Supplementary Figure 4D-H). Given that the up-regulated expression of RAMP2 was variable between mice, we performed a correlation analysis between hepatic RAMP2 levels and AUC following a glucagon challenge test in obese mice but did not observe any correlation (Supplementary Figure 4I). In this experimental setting, therefore, up-regulation of hepatic RAMP2 did not have a dramatic effect on carbohydrate metabolism in adult male mice.

## 4 Discussion

Here, we demonstrate that RAMP2 modulates both trafficking and function of the GCGR. In both HEK293T and Huh7 cells, RAMP2 promotes intracellular accumulation of the GCGR, both basally and following agonist stimulation. Given that FITC-GCG is rapidly deposited intracellularly, presumably by the GCGR, even in the absence of RAMP2 where the GCGR is primarily detected at the plasma membrane, the effect of RAMP2 in GCGR subcellular localisation is likely to be due to a reduction in the rate of GCGR recycling. This conclusion is further supported by our finding that preventing receptor recycling by knocking out the WASH complex similarly leads to intracellular retention of the GCGR in the absence of RAMP2. RAMPs have previously been shown to alter the recycling rate of GPCRs: Bomberger *et al* demonstrated that RAMP3 is responsible for retaining the adrenomedullin receptor 2 (RAMP3/CRLR) intracellularly after agonist stimulation and internalisation [32]. More recently, Mackie *et al* showed that RAMP3 is required for rapid recycling of the atypical chemokine receptor ACKR3 [33]. Taken together, these studies appear to indicate that RAMPs have differential effects on regulating receptor recycling rates, depending on the GPCR in question. We speculated that endosomal GCGR would be more likely to be targeted for degradation over prolonged agonist incubation times. We indeed observed increased lysosomal localisation as well as a trend towards increased degradation of GCGR in the presence of RAMP2 after longer term glucagon exposure, a finding previously made for RAMP3 and the CRLR [32]. Given that RAMPs are ubiquitously present in many tissue types, modulation of their expression levels may play important roles in the control of physiological receptor turnover via fine tuning of the balance between receptor recycling and degradation rates.

The interaction of RAMP2 with GCGR has functional consequences: here we demonstrate that upregulation of RAMP2 acutely increases agonist-stimulated cAMP production in hepatoma cells, a phenomenon that has previously been shown in other non-hepatocyte cell lines [20; 21]. We also observe a similar increase in cAMP production when GCGR is artificially forced to accumulate intracellularly with blocked recycling in WASH-out MEFs, indicating that intracellular GCGR sequestration triggered by RAMP2 co-expression is likely to be responsible for the detected increase in accumulated cAMP. Indeed, using a novel NanoBiT subcellular G protein activation assay we were able to demonstrate that, in the presence of RAMP2, there is greater GCGR activity specifically from endosomes. Data from other secretin-family GPCRs, including the parathyroid hormone receptor and the GLP-1R, which both signal via Gα_s_ from early endosomes [38–40], indicate that the spatiotemporal regulation of signalling is paramount to modulate receptor outputs, with intracellular signalling usually associated with more sustained responses [41; 42]. Our data suggests that a similar phenomenon may be true for the GCGR/RAMP2 complex, a possibility that deserves further investigation especially given its important ramifications for therapeutic targeting of the receptor [43].

We also observed a RAMP2-induced bias away from β-arrestin-2 recruitment at the GCGR, which corroborates our previous findings in CHO cells using a beta-galactosidase fragment complementation assay [20]. β-arrestin recruitment has potentially variable effects on G protein-dependent GPCR signalling: it may terminate G protein-mediated signalling via uncoupling of G proteins and increased activity of cAMP phosphodiesterases [44; 45], but it has also been proposed to facilitate sustained endosomal signalling by the formation of GPCR–G protein–β-arrestin supercomplexes [43]. We have previously demonstrated that, for the GCGR and related GPCRs, absence of β-arrestins increases overall agonist-stimulated cAMP production [26]. In the present study, bias away from β-arrestin-2 recruitment could therefore explain the overall increase in glucagon-stimulated cAMP accumulation when RAMP2 is over-expressed. Since β-arrestins have been shown to be involved in the recycling of GCGR from intracellular compartments back to the plasma membrane, a reduction in their recruitment could be responsible for the increased intracellular retention of GCGR in the presence of RAMP2 [46]. Signalling bias conferred by RAMP2 on the GCGR has previously been reported in a chimeric yeast system, where RAMP2 appeared to reduce recruitment of Gα_i_ and increase recruitment of Gα_s_ to the receptor [21]; however, we observed no differences in recruitment to mini-G_s_, -G_i_ or -G_q_, albeit with very low overall levels of G_i_ recruitment, in our experimental system.

There is very little data in the literature investigating the effects of RAMP up-regulation on GPCR activity *in vivo*. Our *in vitro* findings suggested that RAMP2 may have a role in modifying GCGR activity in hepatocytes, therefore potentially impacting the processes of glycogenolysis, gluconeogenesis and fatty acid oxidation. Here, however, aside from a small improvement in glucose tolerance after a 5hour fast and a trend to greater efficacy for agonist-stimulated cAMP production in isolated hepatocytes, we did not find major effects on carbohydrate metabolism of over-expressing hepatic RAMP2 in either lean or obese mice. It remains to be investigated whether hepatic down-regulation (rather than over-expression) of RAMP2, or RAMP2 up-regulation in other nutritional contexts may yield a biological effect on glucagon signalling *in vivo*. There may also be species-specificity of RAMP2 effects: the *in vitro* data here concerns human RAMP2 interacting with human GCGR, whereas in our *in vivo* experiment we manipulated hepatic murine RAMP2 and therefore studied effects of its interaction with murine GCGR. Furthermore, we cannot rule out an effect of RAMP2 modulation of GCGR in other pathophysiological contexts, such as cirrhosis, which is known to increase RAMP2 expression [47], or in different target organs.

## 5 Conclusion

In the absence of RAMP2, GCGR is found predominantly at the cell surface at steady state, but this localisation is underlaid by rapid and continuous cycles of receptor internalisation and recycling following acute ligand stimulation. In the presence of RAMP2, GCGR accumulates intracellularly, both under basal conditions and following glucagon stimulation. Upon acute stimulation with glucagon, RAMP2 co-expression results in a short-term increase in cAMP accumulation, which may be explained by more efficient signalling from GCGRs retained in endosomes. RAMP2 is also associated with a bias away from β-arrestin-2 recruitment, which may provide a mechanism for the retention of GCGR intracellularly. Over prolonged periods of glucagon stimulation, there is a trend towards increased GCGR degradation, potentially associated with a long-term reduction in efficacy for cAMP production in the presence of RAMP2. Finally, we present *in vivo* data suggesting that these effects on signalling do not lead to a readily observable phenotype in lean and obese mice following up-regulation of hepatic murine RAMP2. Further work is needed to determine the circumstances in which RAMP2 might play a role in regulating GCGR trafficking and signalling *in vivo* in the liver.

## 7 Acknowledgements

We thank David MacDonald for performing the mouse tail vein injections, Eleni Vloumidi for primary hepatocyte harvest and Roger K. Sunahara at University of California, San Diego for helpful discussion on the development of the NanoBiT-Nb37 assay. ERM was supported by a Royal College of Surgeons of England one-year research fellowship and an MRC Clinical Research Training Fellowship while working on this project. AT and DC are funded by the MRC (MR/R010676/1 and MC-A654-5QB10, respectively). AT also acknowledges support from Diabetes UK and the European Federation for the Study of Diabetes. BJ is supported by the MRC, European Federation for the Study of Diabetes, Society for Endocrinology, British Society for Neuroendocrinology, and the NIHR Imperial Biomedical Research Centre. TT and SRB are funded by the NIHR Imperial BRC. AI was funded by the LEAP JP20gm0010004 and the BINDS JP20am0101095 from the Japan Agency for Medical Research and Development (AMED), JST Moonshot Research and Development Program JPMJMS2023 from Japan Science and Technology Agency (JST), and The Uehara Memorial Foundation. The Department of Metabolism, Digestion and Reproduction at Imperial College London, UK, is funded by grants from the MRC and the Biotechnology and Biological Sciences Research Council and is supported by the NIHR Imperial Biomedical Research Centre (BRC) Funding Scheme. The views expressed are those of the authors and not necessarily those of the abovementioned funders, the NHS, the NIHR, or the Department of Health.

## 8 Declaration of Interest

The authors declare no competing interests.

## 10 Supplementary Figure Legends

**Supplementary Figure 1 – Agonist-stimulated internalisation of GCGR *vs*. GLP-1R in the rat pancreatic beta cell line INS-1 832/3 and with GFP-tagged receptors in HEK293T cells.**

A: cAMP dose responses to glucagon (GCG) vs. FITC-GCG in Huh-GCGR cells after 30 minutes stimulation; n = 2; 4-parameter fit of pooled data shown. B: INS-1 832/3 cells transfected with SNAP-GCGR (labelled with SNAP-Surface 549 probe, red) and stimulated with FITC-GCG (green). C: INS-1 832/3 cells transfected with SNAP-GLP-1R (labelled with SNAP-Surface 549 probe, red) and stimulated with FITC-GLP-1 (green). D: HEK293T cells transfected with GCGR-GFP (green) with and without GCG stimulation for 40 minutes. E: HEK293T cells transfected with GLP-1R-GFP (green) with and without GLP-1 stimulation for 40 minutes. Nuclei stained with DAPI (blue); scale bars = 10 μm.

**Supplementary Figure 2 – HALO-tagged GCGR is predominantly localised at the cell surface in the absence of RAMP2 but retained intracellularly in the presence of RAMP2.**

A, B: HEK293T cells transfected with HALO-GCGR (labelled with HALO-AlexaFluor 660 probe, red) with or without co-expressed CLIP-RAMP2 (labelled with CLIP-Surface 547, green) and treated with vehicle (A) or FITC-GCG (grey) (B). Nuclei stained with DAPI (blue); scale bars = 10 μm.

**Supplementary Figure 3 – Agonist-stimulation at the GCGR leads to cAMP production and recruitment of β-arrestin-2.**

A, B: cAMP (A) and β-arrestin-2 (B) dose responses to GCG stimulation in GCGR DiscoverX cells (Eurofins DiscoverX); n = 4; data is mean ± SEM; 4-parameter fit of pooled data shown. C, D: Plasma membrane (C) and endosomal (D) baseline Gα_s_ activity measured by NanoBiT complementation assay in the presence of pcDNA3.1 or RAMP2; n = 5; paired t-test. E: Representative images from Western blots of SNAP-GCGR and tubulin levels in SNAP-GCGR-expressing HEK293T cells with and without RAMP2 co-expression following stimulation with 100 nM GCG for 4 hours; image of membrane cropped as indicated by black surrounding lines and spliced together (see Figure 3L for quantification).

**Supplementary Figure 4 – Up-regulation of hepatic RAMP2 in obese adult male mice is not associated with a change in phenotype.**

A: *Ramp2* gene expression normalised to AAV-alb-GFP, data is mean **±** SEM; n = 4-5 in each group. B: Hepatic RAMP2 protein expression normalised to tubulin; n = 9-10 in each group. C: Western blot depicting RAMP2 (upper panel) and tubulin (lower panel) levels from mice 4 months post-AAV injection; AAV-alb-GFP (GFP) or AAV-alb-RAMP2 (R2) as indicated; mouse brain as positive control. Images of membranes have been cropped and spliced together. D: Body weight. E: Glucose tolerance test. F: Insulin tolerance test. G: Glucagon challenge. H: Pyruvate tolerance test. D-H: Experiments performed in mice 3 to 4 months post-AAV injection, with AUC shown as inset; n = 9-10 per group; data is mean ± SEM. I: AUC for GTT *vs*. protein expression of RAMP2 for each mouse; statistical significance was analysed using 2-way ANOVA with Sidak’s multiple comparison test for time-courses and unpaired t-test for AUCs.

## Notes

### Competing Interest Statement

The authors have declared no competing interest.

