## Supplementary figures and images for "Receptor Activity-Modifying Protein 2 (RAMP2) alters glucagon receptor trafficking in hepatocytes with functional effects on receptor signalling"

### Supp. Fig. 1

#### Supplementary Figure 1

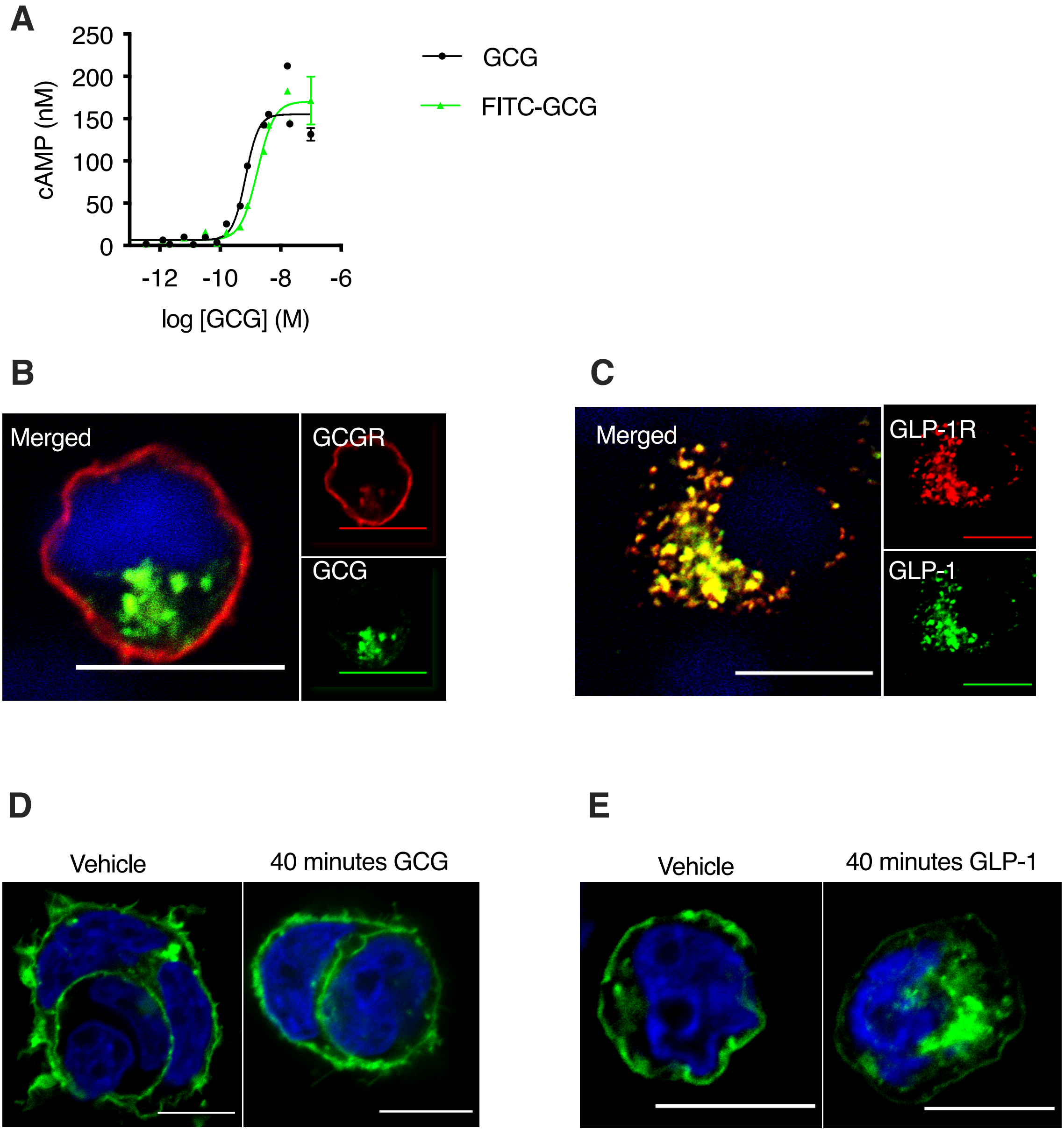

### Supp. Fig. 2

#### Supplementary Figure 2

**A**

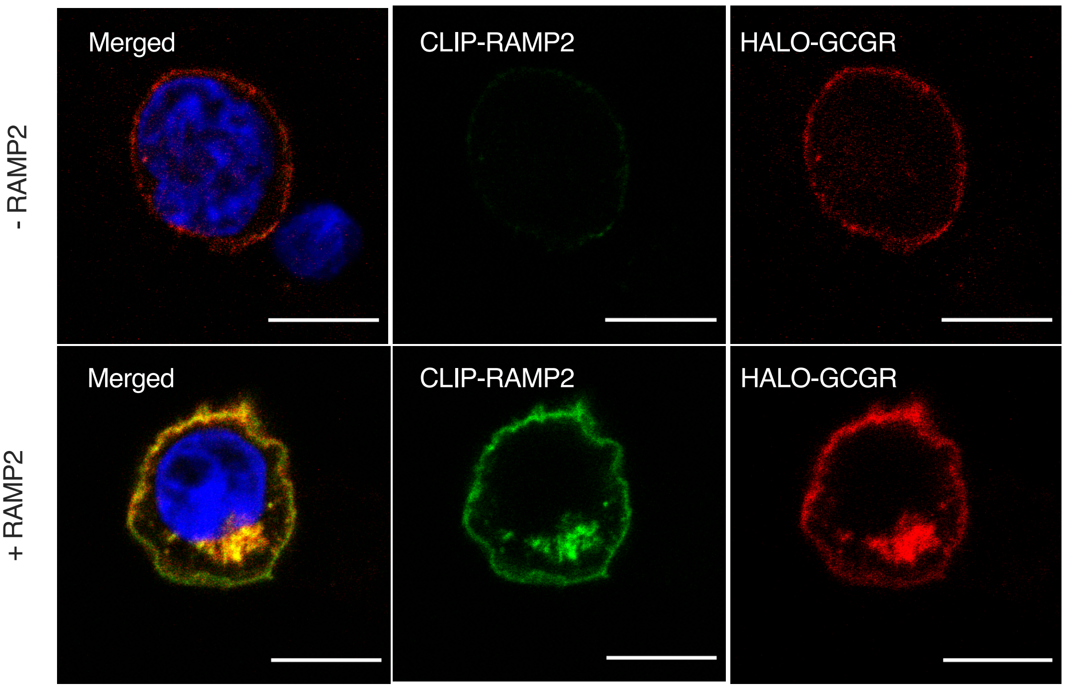


**B**

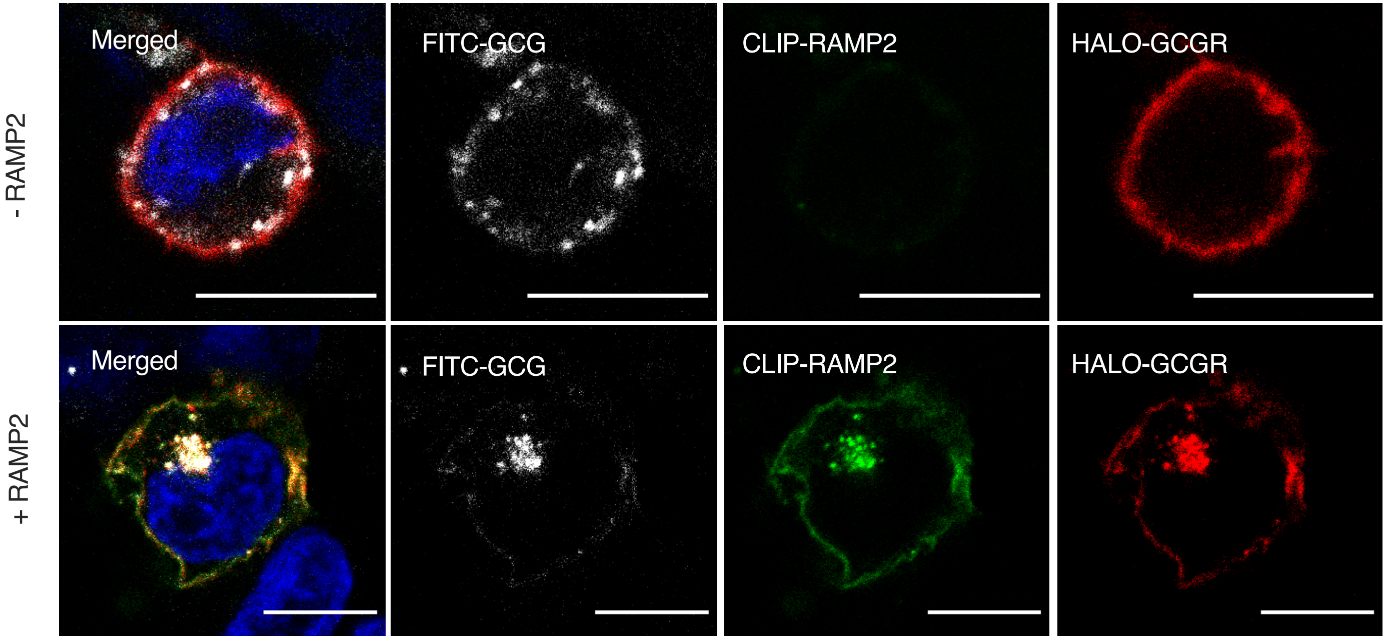

### Supp. Fig. 3

#### Supplementary Figure 3

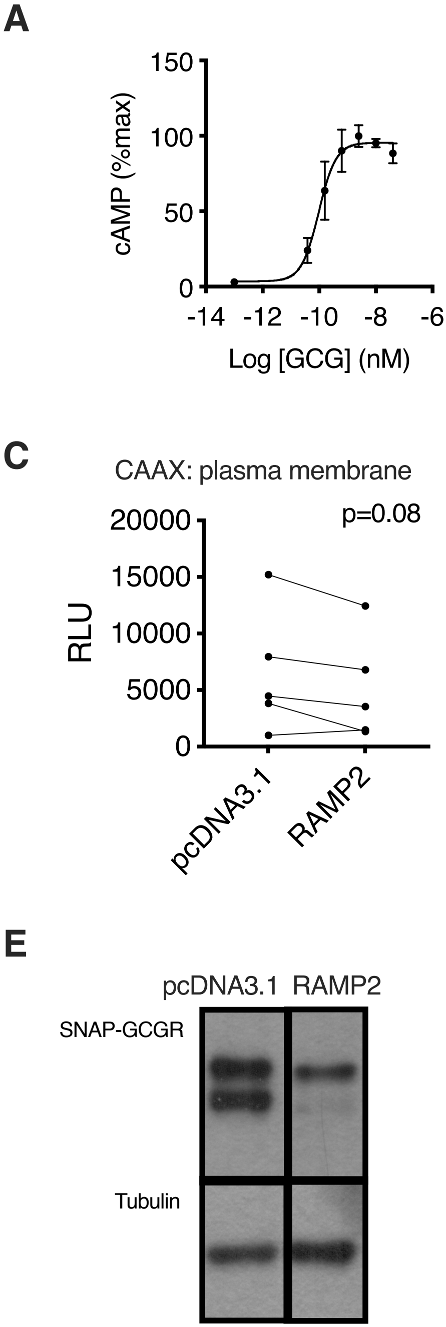

### Supp. Fig. 4

##### Supplementary Figure 4

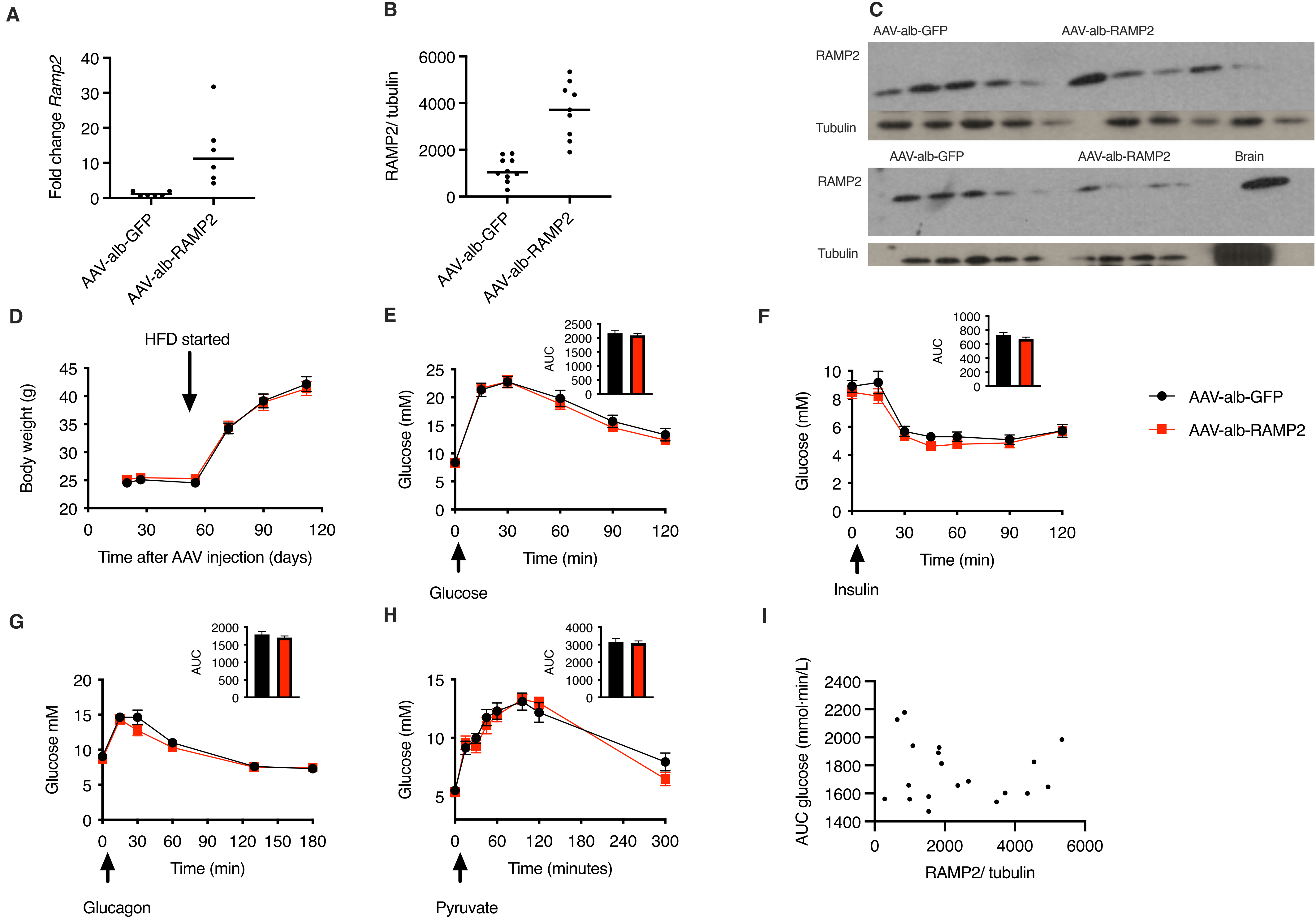
