## Supplementary material for "Receptor Activity-Modifying Protein 2 (RAMP2) alters glucagon receptor trafficking in hepatocytes with functional effects on receptor signalling": Supp. Methods

### Supplementary Methods

#### Labelling and stimulation

Receptor labels (SNAP-Surface 549, SNAP-Surface 649, CLIP-Surface 547 from New England Biolabs; HALO-AlexaFluor 660 from Promega) were applied at a concentration of 1 µM in full media for 30 minutes at 37°C. After washing, cells were stimulated with FITC-peptide in the dark or unlabelled peptide at a concentration of 100 nM in serum free media at 37°C for 30 minutes unless otherwise stated. After treatments, cells were fixed with 4% paraformaldehyde for 30 minutes at 4°C. Coverslips were mounted using mounting media (ProLong diamond antifade, with or without DAPI; Thermo Fisher).

#### Imaging

Images were taken randomly from areas where cells were lying in a single plane using a Zeiss LSM-780 inverted confocal laser-scanning microscope in a 63x/1.4 numerical aperture oil-immersion objective from the Facility for Imaging by Light Microscopy (FILM) at Imperial College London, except from high-content microscopy experiments (Figure 2C and D), which were performed in a an automated Nikon Ti2 widefield microscope with LED light source (CoolLED) and a 0.75 numerical aperture 20X air objective.

*Quantification of Receptor Internalisation:* Confocal microscopy images were analysed in Fiji [1; 2]. In Figure 1, five images were analysed per time point. For each image, 3 plot profiles were drawn through the cell bisecting the cell membrane. Plot profiles were also drawn outside the cells and the average of 3 taken to obtain a background threshold. Plasma membrane boundaries were set by numerically determining the start and end of intensity peaks over background. AUCs were calculated from plot profile histograms after segmentation for the plasma membrane peak, expressed as a percentage of the membrane AUC at time zero, and inversed to calculate percentage of internalisation. Data was non-linearly fitted to an exponential plateau model curve to visualise percentage of receptor internalisation over time. In Figure 2, experiments with Huh7-GCGR cells were performed in 24-mm coverslips, in duplicate. 3 random images were taken per coverslip, and FITC-GCG density calculated for the whole image, subjected to background density subtraction, and normalised to number of cells. For HEK293T cells, quantified data represent mean ± SEM of 8-12 cells for each condition, measured from at least 2 independent experiments. Cells co-expressing the SNAP-GCGR and either EV- or RAMP2-CFP were identified and drawn around by hand. A macro was used to calculate the total and inner cell density of GCGR (as indicated by labelled SNAP-GCGR), the latter calculated after excluding 1 µm around the cell border. Density of receptor inside the cell was expressed as a ratio of its density across the total cell. Statistical comparisons were performed using unpaired t-tests.

#### Radioligand whole cell binding assays

Cells in suspension were incubated with I^125^-glucagon in assay buffer (25mM HEPES pH 7.4, 2 mM magnesium chloride hexahydrate powder, 0.05% Tween 20, 1% bovine serum albumin (BSA) and 0.5% protease inhibitor; Sigma), 48 hours after transfection, as previously described [3]. All incubations were performed at 4°C for 4 hours, which is sufficient to ensure binding equilibrium [4]. Non-specific binding was determined by adding an excess of non-labelled glucagon (3 µM) for each concentration of I^125^-glucagon. After washing, pellets were measured for gamma radiation for 240 seconds (Gamma Counter NE1600, NE Technology, UK). Specific binding was calculated by subtracting non-specific from total binding at each concentration. All conditions were tested at least in triplicate.

#### cAMP assays

All assays were performed at 37°C. For MEF cell experiments, cells were resuspended in serum-free DMEM with phosphodiesterase inhibitors (IBMX 100 µM) and glucagon at indicated concentrations for 10 minutes before lysis and application of cAMP detection reagents as per manufacturer's instructions (cAMP Dynamic 2, Cisbio). For Huh7-GCGR and primary hepatocytes, assays were performed on plated cells, to which glucagon was added at the indicated concentrations for the indicated period before lysis. Four-parameter curve fitting was performed using Prism 8.0 (GraphPad Software).

#### NanoBiT complementation assays

All nanoBiT assays were performed in HEK293T cells.

*Mini-G protein/β-arrestin-2 recruitment assays:* The GCGR-SmBiT plasmid was generated by in-frame cloning of the SmBiT tag (SmBiT F: 5′-ccggtggtggatccggcggaggtgtgaccggctaccggctgttcgaggagattctgtaat-3′; SmBiT R: 5′-gatctaatgtcttagaggagcttgtcggccatcggccagtgtggaggcggcctaggtggt-3′) at the C-terminus of GCGR by substitution of the Tango sequence on a FLAG-tagged GCGR-Tango expression vector [5] (a gift from Prof Bryan Roth, University of North Carolina, USA; Addgene plasmid #66291). Mini-G_s_, mini-G_q_, and mini-G_i_ plasmids, tagged at the N-terminus with the LgBiT tag [6], were a gift from Prof Nevin Lambert, Medical College of Georgia, USA. For β-arrestin-2 recruitment assays, β-arrestin-2 fused at the N-terminus to LgBiT (LgBiT-β-arrestin-2; Promega, plasmid no. CS1603B118) was the configuration chosen, as it has previously been used successfully with another class B GPCR [7]. Cells were seeded in 12-well plates and co-transfected with 0.05 μg each of GCGR-SmBiT and either LgBiT-mini-G_s_, -mini-G_q_, -mini-G_i_ or -β-arrestin-2; plus 0.5 μg of pcDNA3.1 or RAMP2.

*Nb37 bystander assays:* Nb37 (gene synthesised by GenScript with codon optimisation) was C-terminally fused to SmBiT with a 15-amino-acid flexible linker (GGSGGGGSGGSSSGGG), and the resulting construct was referred to as Nb37-SmBiT. The C-terminal KRAS CAAX motif (SSSGGGKKKKKKSKTKCVIM) was N-terminally fused with LgBiT (LgBiT-CAAX). The Endofin FYVE domain (amino-acid region Gln739-Lys806) was C-terminally fused with LgBiT (Endofin-LgBiT). Gαs (human, short isoform), Gβ1 (human), Gγ2 (human), and RIC8B (human, isoform2) plasmids were produced in house. These constructs were inserted into the pcDNA3.1 or the pCAGGS expression plasmid vectors. Cells were seeded in 6-well plates and co-transfected with 0.2 μg SNAP-GCGR, 0.5 μg Gαs, Gβ1, and Gγ2, 0.1 μg RIC8B, 0.1 μg CAAX-LgBiT or 0.5 μg Endofin-LgBiT with 0.1 μg or 0.5 μg Nb37-SmBiT, respectively, plus 0.2 μg RAMP2 or pcDNA3.1.

Briefly, 24 hours after transfection cells were detached, resuspended in NanoGlo Live Cell Reagent (Promega) with furimazine (1:20 dilution) and seeded into white 96-well half area plates. Baseline luminescence was recorded over 5 minutes at 37°C in a Flexstation 3 plate reader, and for 30 minutes with or without addition of glucagon at 100 nM for G protein and β-arrestin-2 recruitment assays, and at serial doses of up to 1 μM for the Nb37 bystander assays; readings were taken every 30 seconds or every minute, respectively. Readings were normalised to well baseline and then to average vehicle-induced signal to establish the agonist-induced effect. AUCs from response curves were calculated for each concentration of glucagon and fitted to four-parameter curves using Prism 8.0 (GraphPad Software).

#### Degradation assay

HEK293T cells in 6-well plates were co-transfected with 1.5 µg each of SNAP-GCGR and either human RAMP2 or pcDNA3.1. The cells were detached 24 hours later, split into two and allowed to re-adhere in 12-well plates for 4-6 hours. The cells were then washed and stimulated with vehicle or 100 nM glucagon in serum-free medium overnight. Lysates were collected the next day by adding 1X TNE lysis buffer (20 mM Tris, 150 mM NaCl, 1 mM EDTA, 1% NP40, protease and phosphatase inhibitors) for 10 minutes at 4°C followed by cell scraping and sonication (3X, 10 seconds each). The lysates were then frozen at -80°C for 2 minutes, thawed and centrifuged for 10 minutes at 4°C and 15,000 × g. The supernatants were collected, fractionated by SDS-PAGE in urea loading buffer and analysed by Western blot. Blots were developed with the Clarity Western enhanced chemiluminescence (ECL) substrate system (BioRad) in a Xograph Compact X5 processor and specific band densities quantified in Fiji.

#### Analysis of gene and protein expression in mouse liver

RNA extraction was performed using RNeasy mini kit (Qiagen), reverse transcription using High-Capacity cDNA Reverse Transcription kit (Thermo Fisher) and quantitative PCR using the Taqman^TM^ system (with probe Mm00490256_g1 for *Ramp2* and Hs99999901_s1 for endogenous control gene 18S), all according to manufacturer’s instructions.

For protein extraction, snap-frozen liver tissue was lysed in ice-cold RIPA lysis buffer (Sigma) containing protease inhibitor cocktail (Promega) using a TissueLyser machine (Qiagen). Samples were agitated for 2 hours at 4°C to allow proteins to dissolve in the buffer. Next, samples were centrifuged at 12,000 × g at 4°C for 10 minutes, and the insoluble pellet discarded. Protein content of the soluble fraction was quantified using bicinchoninic acid assay kit (Sigma). 30 μg of protein per sample were resolved using SDS-PAGE and analysed by Western blot as above.
